# A universal pipeline for the alignment of electrode tracks with slice histology and electrophysiological data

**DOI:** 10.1101/2022.06.20.496782

**Authors:** Jorrit S. Montijn, J. Alexander Heimel

## Abstract

Modern electrophysiological experiments in neuroscience typically measure the activity of hundreds of neurons across multiple brain regions. Each recorded neuron has to be assigned its correct area of origin, which can be estimated using histological reconstructions of electrode tracks. However, this approach has a limited accuracy with which the probe position can be determined. While some tools exist to assist in this process in mice, such tools are scarce for other species, such as rats or macaques. Moreover, even in mice, a more reliable brain area assignment may be achieved by using spiking response properties to delineate area boundaries along the recording probe. Here we present the Universal Probe Finder, which provides multi-species support for reconstructing a probe’s brain area per spike-sorted cluster or recording site. It can use various data formats and can calculate neuronal responsiveness to experimental interventions with only event times and spike times to provide further neurophysiological markers that simplify and improve the reliability of assigning brain areas to neurons. The program can be downloaded here: https://github.com/JorritMontijn/UniversalProbeFinder.

## Introduction

Modern silicon probes, such as Neuropixels, typically yield hundreds of neurons per recording session across multiple brain areas (Jun et al. 2017; Steinmetz et al. 2019). This has greatly accelerated the pace of neuroscience research, and allows new questions to be tackled (Engel and Steinmetz 2019). While advantageous in many regards, data obtained with high-density silicon probes also require much more data pre-processing. Some steps, such as spike detection and clustering, can be mostly automated (Pachitariu et al. 2016). Others, however, like histological reconstructions of the probe location, are still performed mostly manually (Shamash et al. 2018). Especially extracting brain area locations for each spike-sorted cluster is a laborious and error-prone step, since many neurons must be linked to their areas, with multiple areas per probe.

Electrophysiological properties may be used to improve the determination of area boundaries along the length of a high-density silicon probe. For example, multi-unit activity is often highly correlated for contact points outside the brain, and spiking rates are often higher in the hippocampal formation than in subcortical structures. Some excellent tools exist that use these properties (Shamash et al. 2018; Peters [2019] 2022), but – to the best of our knowledge – none use differences in neuronal stimulus-responsiveness as a marker for area boundaries. When optogenetically stimulating a specific region, cells in that area will respond strongly, while areas that do not receive direct stimulation do not show a spiking response (Packer et al. 2012). The same holds for visual cortical and subcortical areas responding to visual stimuli, or auditory areas responding to auditory stimuli, etc. (Chen et al. 2018). These markers can be therefore be extremely informative when delineating area boundaries, but are rarely used. To conclude, current tools are very useful, but have several shortcomings: they are often highly data format-specific, do not use differences in neuronal stimulus-responsiveness as a marker for area boundaries, and/or support only a single species, such as mice.

Here we present the Universal Probe Finder, a histology processing and probe alignment pipeline that uses the recently developed zeta-test to calculate neuronal responsiveness based only on single-unit or multi-unit spike-times and event onsets (Montijn et al. 2021). The Universal Probe Finder supports atlases from multiple species, such as mouse, rat, and macaque, and can use various electrophysiological data formats, including kilosort output and raw SpikeGLX files (Pachitariu et al. 2016; billkarsh [2016] 2022; Allen Institute for Brain Science 2017). To compute neuronal responsiveness, it uses pipeline-independent stimulation files that need only an array of event times. We expect the versatility and extendibility of this pipeline will make it a useful tool for neuroscientists from a wide array of backgrounds.

## Methods

### General workflow

The Universal Probe Finder consists of three integrated programs that allow users to annotate electrode tracks in the Slice Prepper, align slices to an atlas in the Slice Finder, and finally fine-tune the position of the probe in the Probe Finder (fig. 1). Moreover, the Probe Finder can import other probe location formats, such as those from SHARP-track and AP_Histology to further process previously obtained coordinate estimates. The Universal Probe Finder can import various formats, and is built to minimize the work required to add new electrophysiological data formats or brain atlases. Currently the Probe Finder can import Kilosort, SpikeGLX and Acquipix data formats, and use mouse, rat and macaque brain atlases. If annotated and template volumes are available, adding an atlas should take under an hour. All code is available on github (Montijn [2022] 2022).

**Figure 1.**
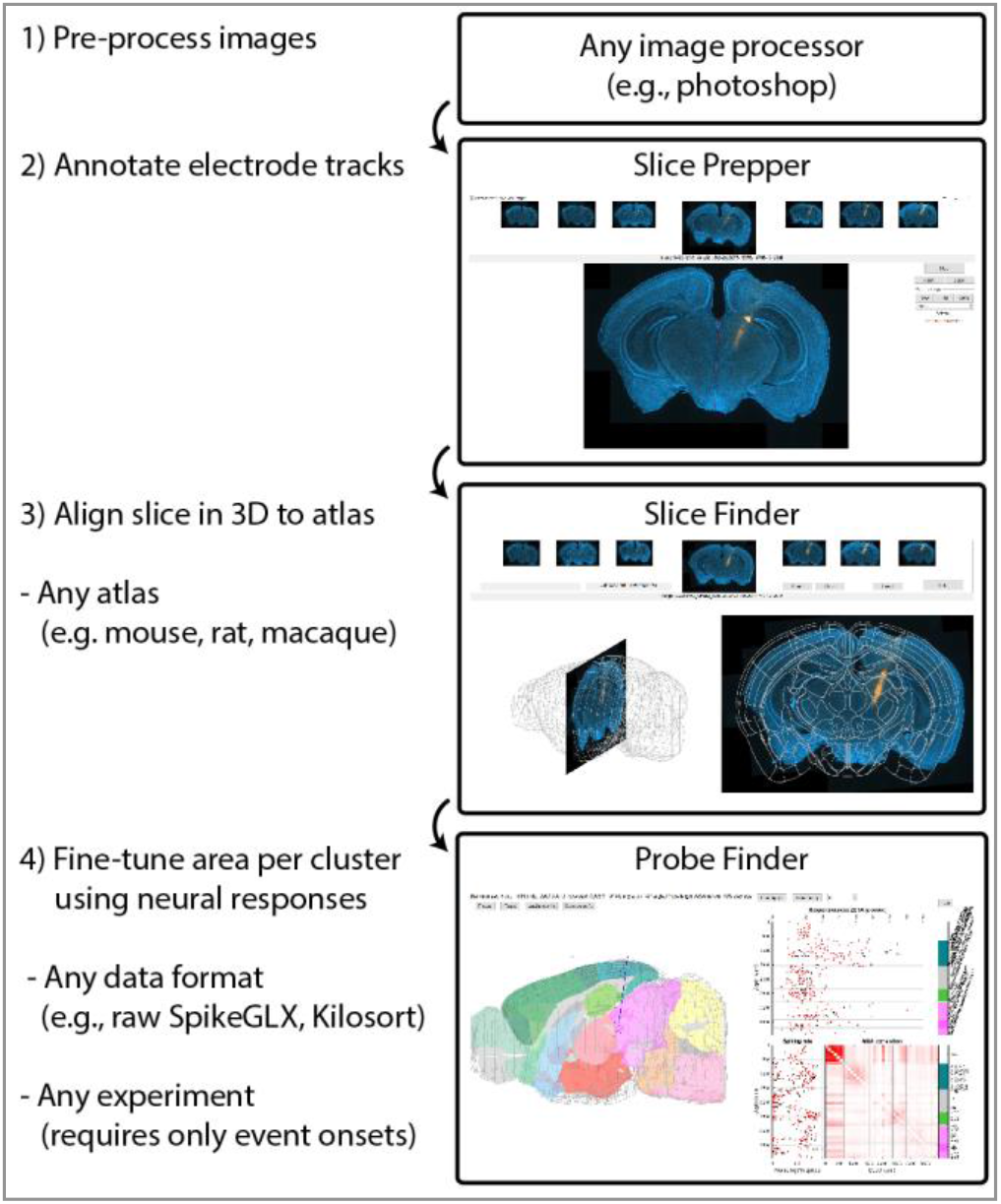
Workflow overview of the Universal Probe Finder. Pre-processed images can be annotated with electrode tack locations, aligned to a 3D atlas, and annotated track locations can then be used as a starting point to further fine-tune the probe location and area boundaries using neural responses.

### Definition of ML/AP angles in the Probe Finder

Rotations along a single axis in R^3^ define a 2D plane, where each point on this plane is defined by a unique combination of an angle and a distance to the axis. Unfortunately, combining two 360-degree axes of rotation in 3-dimensional space leads to overlapping projections, where multiple combinations of two angles can describe the same point. Spherical coordinate systems solve this problem by restricting one of these axes to 180 degrees: i.e., the elevation. For our purposes of defining a probe’s location, it is most useful to have two axes that align more or less with our two most important axes: ML (medial-lateral) and AP (anterior-posterior). Moreover, a natural angular “origin” for the probe would be to define the direction going straight down along the DV (dorsal-ventral) axis as 0 degree ML and 0 degree AP. This requires rotating and flipping the spherical coordinate system, such that the origin is the probe tip, the elevation is the AP angle, and the azimuth is the ML angle (fig. 2).

**Figure 2.**
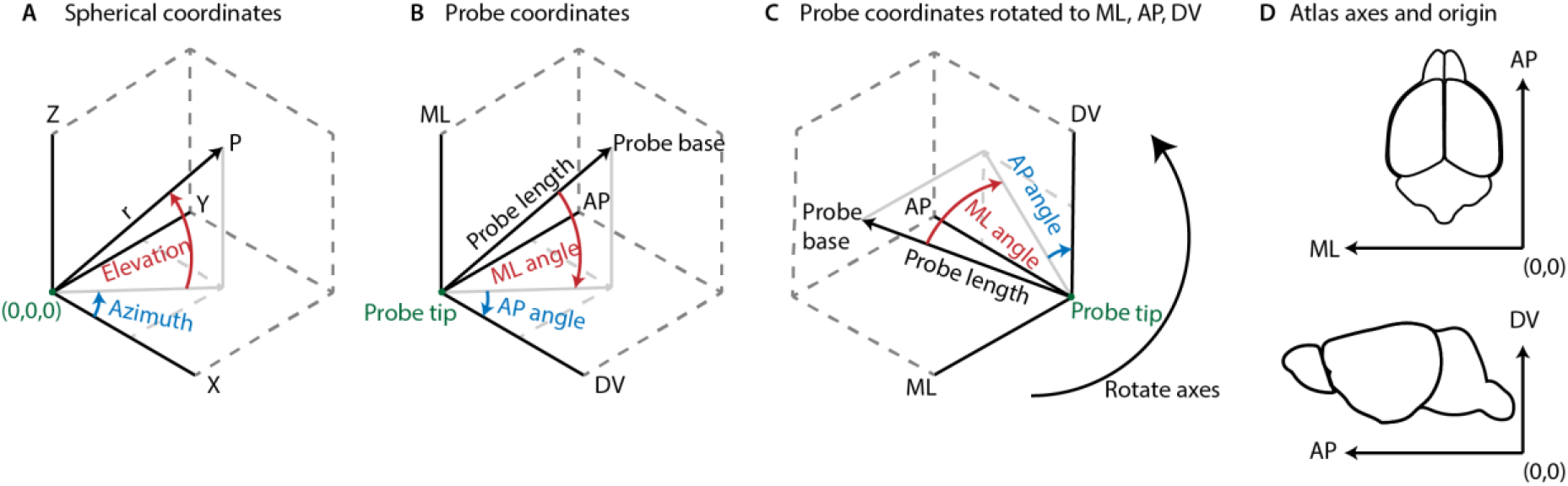
Definition of ML and AP angles. A) Standard spherical coordinate system using elevation and azimuth. Azimuth is defined on the interval [-π, π] and elevation on [-π/2, π/2]. B) X,Y,Z, replaced by stereotactic axes. Note the change in angle direction. C) Rotated axes aligned with atlas directions. D). Example showing the directions of the native atlas axes: (0,0,0) is at the right, bottom, posterior end of the atlas space. Note “origin-centered” coordinates refer to coordinates that are centered relative to, for example, bregma. These origin-centered coordinates flip the ML axis, where right is positive, and left is negative.

### The atlas metadata format

The Universal Probe Finder reads a configuration .ini file to find which atlases are installed. The configuration file specifies the name of the atlas, the name of path variable, the name of the loader function that pre-processes the atlas files, and the amount of downsampling when plotting an atlas slice. The loader function reads the atlas files at the specified path and outputs a structure sAtlas with various fields (table 3).

### The electrophysiology data format

Similar to when loading an atlas, the Universal Probe Finder reads a configuration .ini file to find which electrophysiology formats are installed. An *ephys loader* function receives a directory as input, and outputs various processed electrophysiology variables that are read from that directory (table 4).

### Format of the Probe Finder input file

The initial probe location is calculated from a set of points in atlas-space. In order for the Probe Finder to read a file successfully, the file should contain the variable *sProbeCoords* with at least the field *cellPoints*: a [1 x N] cell array where each cell contains points for a single probe *i*. Here, each entry is a [P x 3] array with P points in [ML AP DV] native-atlas format coordinates. It also possible to add an optional field *cellNames*, a [1 x N] cell array where each cell contains the probe’s name. Note that the Slice Prepper and Slice Finder automatically generate these files when exporting data.

### Format of the Probe Finder output file

The probe location at the moment of data export is saved in the field *sProbeAdjusted* of the structure *sProbeCoords*. The *sProbeAdjusted* structure contains the various fields that describe the location of the probe in atlas-space as well as relative to the origin. Moreover, if electrophysiological data was loaded, it also contains the brain area for each channel and spike cluster (table 1, table 2).

**Table 1.** Probe Finder output file variables. Class is double, unless noted otherwise.

| Name: | Size: | Content |
| --- | --- | --- |
| probe_vector_cart | [2 x 3] | cartesian tip and base coordinates in [ML,AP,DV] atlas space (see fig. 2D) |
| probe_vector_intersect | [1 x 3] | coordinates of brain entry in atlas space |
| probe_vector_sph | [1 x 6] | spherical coordinates in atlas space:<br>[ML AP DV deg-ML deg-AP length] |
| probe_vector_bregma | [1 x 6] | as above, but origin-subtracted |
| probe_area_ids_per_depth | [1 x N] | vector of area IDs per point along the probe |
| probe_area_labels_per_depth | [N x 1] | acronyms per point, cell array |
| probe_area_full_per_depth | [N x 1] | full names per point, cell array |
| probe_area_boundaries | [1 x P+1] | locations along the probe of area boundaries |
| probe_area_centers | [1 x P] | centers of areas |
| probe_area_ids | [1 x P] | id per area of the above |
| probe_area_labels | [P x 1] | acronym per area, cell array |
| probe_area_full | [P x 1] | full name per area, cell array |
| probe_area_ids_per_cluster | [C x 1] | area IDs per cluster |
| probe_area_labels_per_cluster | [C x 1] | area acronyms per cluster, cell array |
| probe_area_full_per_cluster | [C x 1] | full name of the area per cluster, cell array |
| stereo_coordinates | Table | probe angles/coordinates of brain entry in $\mu\text{m}$ relative to the atlas origin; see table 2 |

**Table 2.** Contents of the “stereo_coordinates” variable.

| **Name:** | **Unit** | **Content:** |
| --- | --- | --- |
| ML | Microns | Probe’s brain entry location along ML (medial-lateral) axis, relative to the atlas origin (e.g., bregma) |
| AP | Microns | Probe’s brain entry location along AP (anterior-posterior) axis, relative to the atlas origin (e.g., bregma) |
| AngleML | Degrees | Angle of the probe in the ML direction |
| AngleAP | Degrees | Angle of the probe in the AP direction |
| Depth | Microns | Depth below the brain entry of the lowest recording channel of the probe (i.e., usually the depth of the tip) |
| ProbeLength | Microns | Length of the probe after stretching/shrinking |

**Table 3.** Variables returned by an atlas loader function.

| Name | Size | Description |
| --- | --- | --- |
| av | [ML x AP x DV] | Annotated volume, where a value in *av* denotes an area ID that can directly index into *st*. |
| tv | [ML x AP x DV] | Grey-scale template volume in range [0 255] |
| st | [N x T] table | N-entry table, where N is highest area index in *av*. It must contain *id*, *name*, *acronym* and *parent_structure_id*. |
| Bregma | [1 x 3] | Reference point of atlas in native atlas coordinates, such as bregma, in [ML AP DV] voxel coordinates |
| VoxelSize | [1 x 3] | Size of a single voxel in microns [ML AP DV] |
| BrainMesh | [P x 3] | Mesh of brain outline, where each vertex is a [1 x 3] point, and curve-ends are denoted by a [*nan nan nan*] entry. |
| Colormap | [N x 3] | N-entry RGB color map, specifying a color for each area |
| Type | string | Name, e.g.: *Allen-CCF-Mouse* |

**Table 4.** The *ephys loader* function outputs a structure with the variables below.

| Name | Size | Description |
| --- | --- | --- |
| dblProbeLength | [1 x 1] | Length of the probe in microns |
| vecUseClusters | [1 x N] | Cluster id, where N is the number of used clusters/channels |
| vecNormSpikeCounts | [1 x N] | Normalized spike counts per cluster |
| vecDepth | [1 x N] | Depth below base of probe per cluster/channel |
| vecZeta | [1 x N] | Zeta-responsiveness values (or contamination) |
| strZetaTit | [string] | If no zeta values are present, this is 'Contamination (%)' |
| cellSpikes | [1 x N] | Spike times per cluster/channel |
| ClustQualLabel | [1 x N] | Label per cluster (e.g.: “mua”, “good”) |
| ContamP | [1 x N] | Contamination percentage per cluster |

## Results

All code is available on github: https://github.com/JorritMontijn/UniversalProbeFinder. For users who do not have a MATLAB license, we have also provided a stand-alone installer that does not require a license.

### Slice Prepper

The first step in the Universal Probe Finder pipeline is annotating electrode tracks with the Slice Prepper (fig. 3). The Slice Prepper can read various image formats and prepares all images selected by the user by downscaling and rotating the image. The user can then draw multiple tracks or segments on various slices, and export the pre-processed images and their track annotations for further use in the Slice Finder.

**Figure 3.**
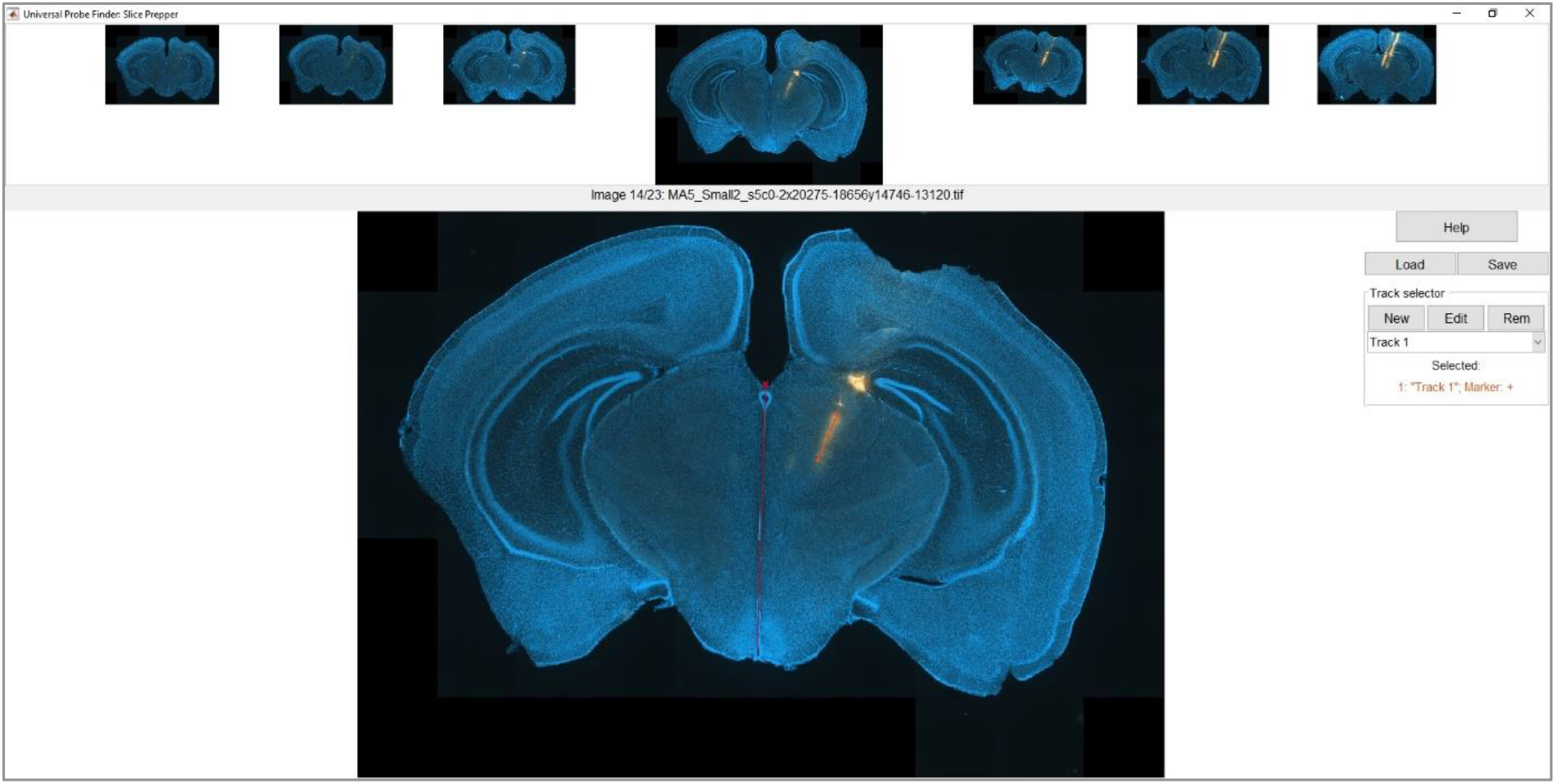
Example view of the Slice Prepper, the first step in the Universal Probe Finder pipeline. The Slice Prepper allows users to prepare their image slices by setting the image ordering, downscaling, aligning the midline, and annotating each slice with electrode tracks. Each track can have a manually specified name, colour and marker for easy referencing.

### Slice Finder

The slices that were annotated with the Slice Prepper can be loaded in the Slice Finder, where each slice can be compared to the brain atlas the user has chosen. The Slice Finder allows each slice to be independently rotated, resized, and translated in medial/lateral (ML), anterior/posterior (AP), and dorsal/ventral (DV) directions (fig. 4). It shows a side-by-side view of the slice location in the brain in 3D, and the annotated slice plus an overlay of the boundaries of the brain atlas for the current slice’s location in the brain. Controls can be switched between atlas-centered and slice-centered to allow accurate brain area boundary alignment to the histology slice.

**Figure 4.**
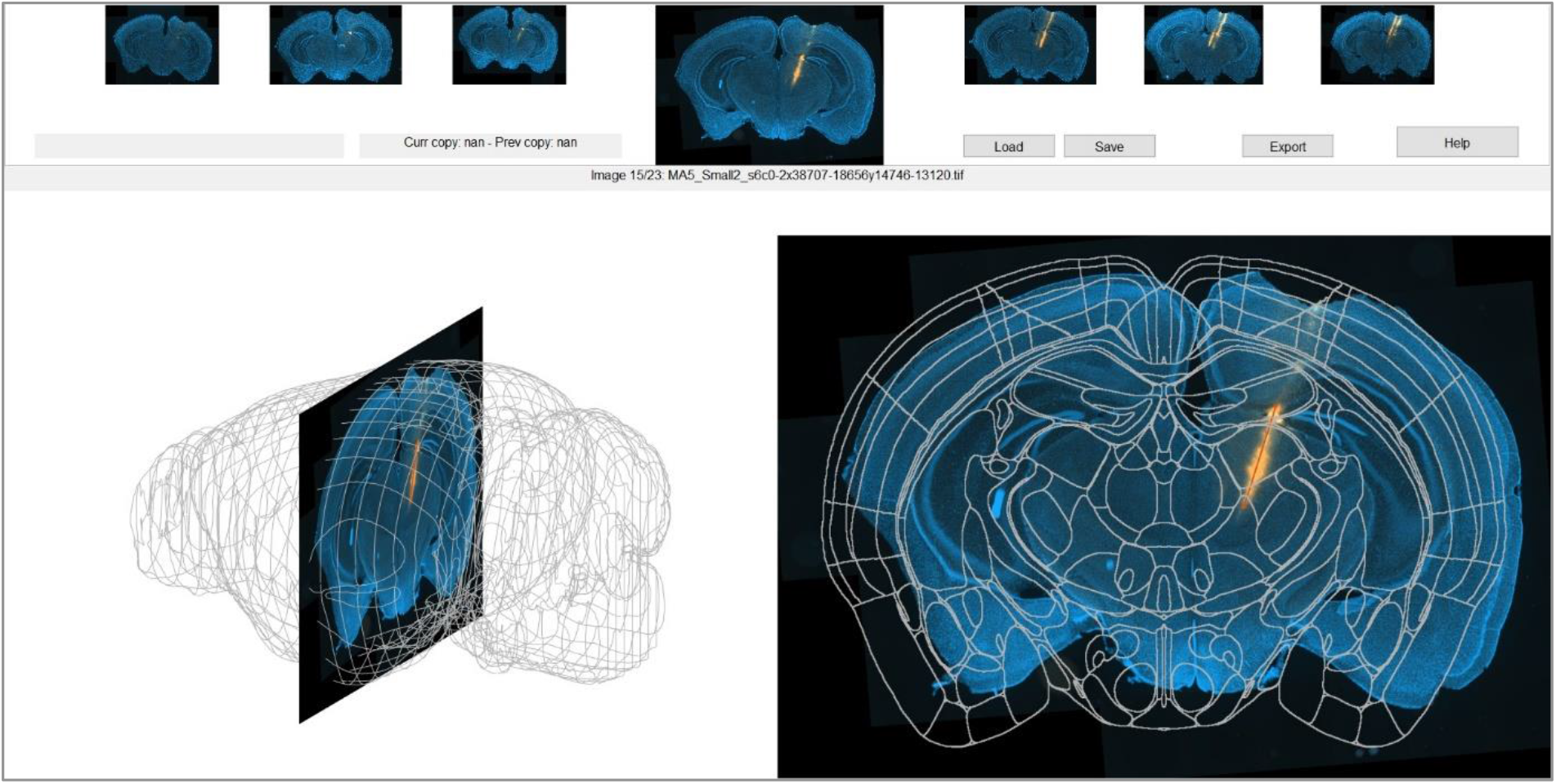
The Slice Finder shows a side-by-side view of the slice in 3D space in the brain (left) and the annotated slice with atlas area boundaries overlaid (right). Position and rotation interpolation across image slices allow high-throughput processing by minimizing the number of images that need manual aligning.

An important addition to the Slice Finder compared to other alignment programs is the ability to copy and paste slice locations between images; and to interpolate the position of intermediate slices between two reference slices. For example, one could align a slice at the anterior end of the set of slice images and one at the end, and then interpolate all other slices. Assuming the slicing angle is the same for all images, and slice thickness is constant, then one needs only to align two slices manually and apply the interpolation for all slice images to correctly register their locations to the brain atlas.

### Probe Finder

After aligning slice locations with the Slice Finder, one can import the positions of the annotated tracks in atlas-space coordinates into the Probe Finder (fig. 5). It calculates the first principal component from all points that make up the start and end points of all annotated track segments. This direction is used as the initial probe vector, and it sets the initial position such that the probe base is exactly at the brain intersection. The brain areas in which electrode channels and spike clusters were located, was historically determined solely on the basis of these histological reconstructions. While this gives an approximate estimate, this estimate is rather coarse, especially when assigning various areas along the probe, as is often the case for modern probes with high recording site density.

**Figure 5.**
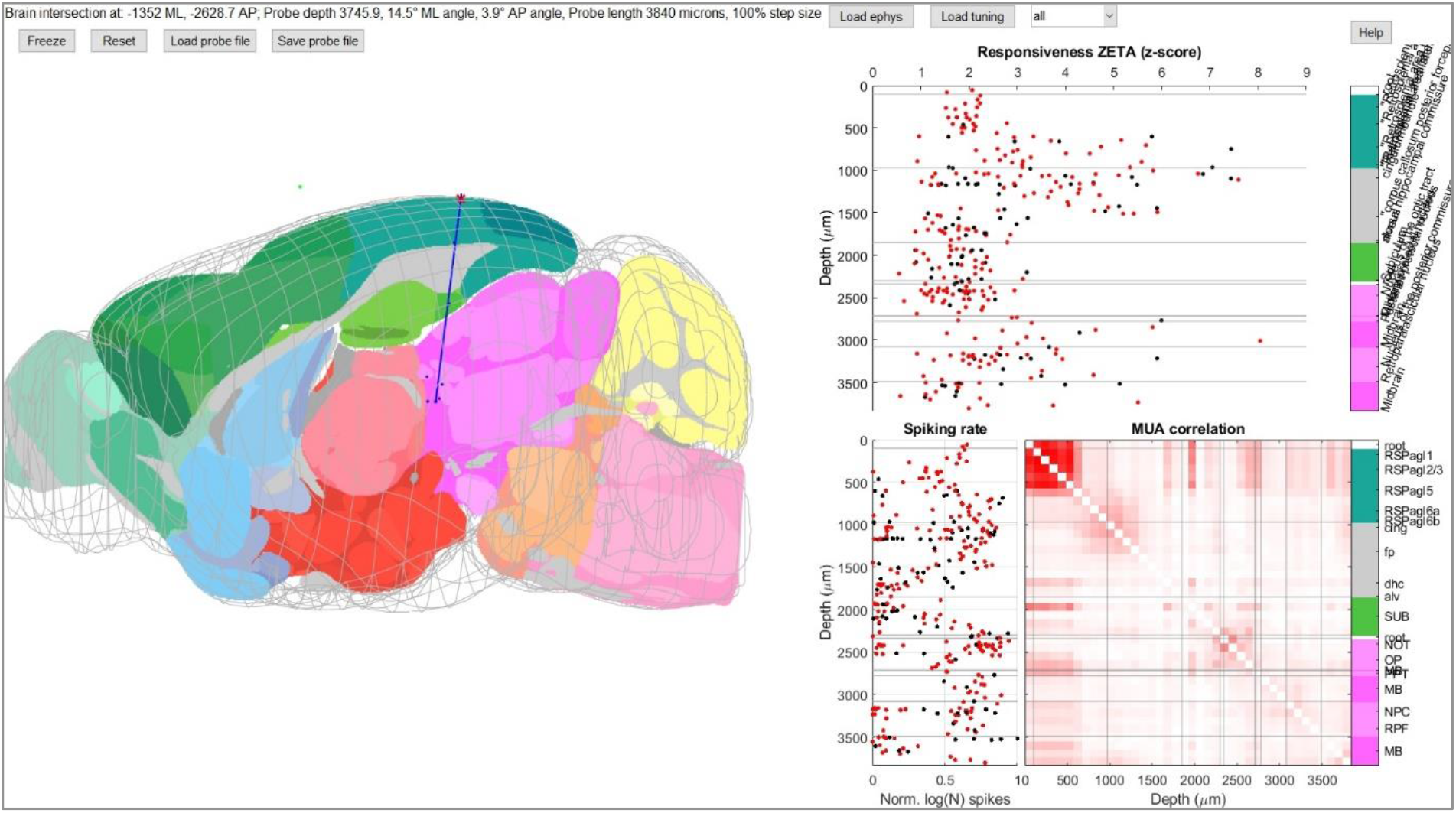
The Probe Finder shows a side-by-side view of the probe in 3D space in the brain (left) and neural response markers per spike cluster or per electrode channel (right). It displays three markers: 1) ZETA-test responsiveness z-scores (top), indicating a neuronal response to the experiment; 2) normalized spiking rates (bottom left); and 3) multi-unit activity (MUA) correlations per set of electrode channels (bottom right). Note that the initial position based on histological data, as shown here, does not match the area boundaries indicated by neural response markers.

The Probe Finder therefore calculates multiple neuronal response markers that allow a more reliable alignment of the probe’s recording sites to the atlas’s brain areas. The neuronal response markers can be calculated at a single recording channel basis, or on single units obtained with spike sorting programs. The Probe Finder calculates three complementary metrics that provide different types of insight into where area boundaries might occur along the recording probe (fig. 6): 1) A normalized spiking rate for each recording location gives a strong indication of the presence of white matter bundles; 2) Multi-unit activity (MUA; here aggregate spike counts over nearby clusters/channels) correlation at a 10 ms time window is strong for noisy contact points not inside the brain; 3) Responsiveness ZETA z-scores shows a clear delineation between areas that respond to the experiment versus those that do not (Montijn et al. 2021). An example experiment with visual stimuli creates a sharp boundary in responsiveness values between visually-responsive and non-visually responsive regions (fig. 6). Using all three markers to align the probe can therefore significantly improve the reliability of area assignments over histology-only alignments.

**Figure 6.**
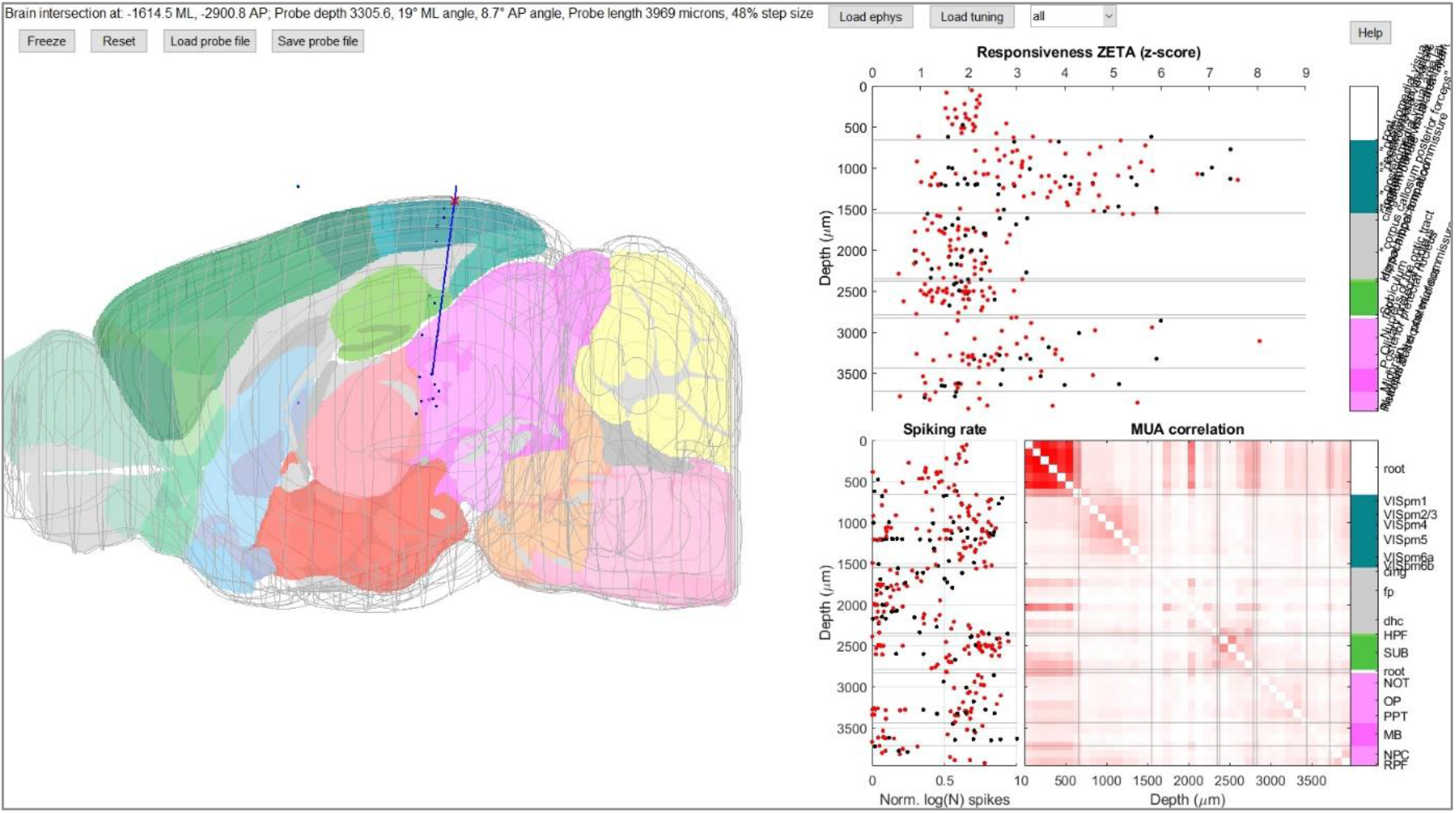
A view of same experiment as figure 5, but with better area alignments. The three neuronal response markers shown in the Probe Finder can discriminate different types of areas: 1) normalized spiking rates are low in white-matter bundles (here, fp) and high in brain areas; 2) multi-unit activity (MUA) correlations are provide a sharp boundary between electrodes outside the brain (high correlations due to noise) and inside the brain (lower correlations). The MUA correlations may also help to distinguish between areas, as correlations are generally higher within areas than between areas. 3) ZETA-test responsiveness z-scores can provide very sharp delineations between areas of interest that respond to the experiment, and those that do not (here for example VISp versus white matter bundles; and visual midbrain areas versus subiculum).

## Discussion

The Universal Probe Finder provides a processing pipeline for histology slices and probe alignment that requires no programming or code editing by its users. It can also be used to explore various brain atlases for planning experiments. Compared to other processing packages, the Universal Probe Finder can use a wider range of brain atlases, read multiple electrophysiological data formats, and compute neuronal responsivity to the experimental stimuli. Moreover, the ability to interpolate the rotation and position of slices to intermediate images means that users only need to perform manual slice alignment for two images per experiment. The Universal Probe Finder is open source and easily modifiable, so users can add custom atlases and data formats. Finally, users who do not possess a MATLAB license can install a stand-alone package.

While the Universal Probe Finder improves upon other packages in various ways, probe position reconstructions are still subject to some mislocalization errors. One of the most important causes of poor reconstruction is tissue warping during brain slicing and when mounting the slices on a cover glass. Avoiding brain slicing and mounting altogether will therefore considerably reduce these problems. Histological reconstructions may be improved by instead performing 3D whole-brain imaging of clarified tissue, but this will require more specialized skills than slicing and mounting (Chung and Deisseroth 2013). As long as probe location reconstructions are based on image slices, the location error can be rather large when using only histological data. While these inaccuracies are still present in the Universal Probe Finder pipeline, its impact on the final area assignment per cluster/channel is considerably reduced because the alignment step uses neural spiking activity as markers for area boundaries. Even if histology will be performed on 3D whole-brain images, area assignment accuracy will likely still benefit from using neurophysiological markers, such as zeta-responsiveness, spiking rates, and MUA correlations.

## Acknowledgements

We thank Nora Jamann, Chris Eva, Frederic Michon, Chris Klink, and Robin Haak for beta-testing. We thank Andy Peters and Philip Shamash for inspiration on the design. Jorrit Montijn was funded by the Fonds KNAW-instituten.

